# LongVerse: verified AI agents for Nanopore and PacBio DNA methylation analysis

**DOI:** 10.64898/2026.10.13.758965

**Authors:** Yang Liu, Qiuyang Zhang, Sheng Li

## Abstract

AI agents now run sequencing pipelines, but their outputs are checked by rubrics or language-model judges, not against an independent measurement. LongVerse turns one English sentence into haplotype-resolved DNA methylation from Nanopore or PacBio reads, through agents that read the data, supply arguments to a pinned pipeline and never write code. On HG002, a healthy reference genome, agent and human expert produced byte-identical calls (r = 1.000) that agreed with bisulfite sequencing (r = 0.943, 0.884), at $0.12 to $0.54 of conversation per run. A consensus agent that admits any per-site caller reduced error 10.7 percent against the best published caller.

## Main

AI agents now operate bioinformatics pipelines, and they differ in what they verify (**Extended Data Table 1**). Paper2Agent turns a paper and its repository into a Model Context Protocol (MCP) server and is graded against human-executed answers, on tools that finish on a laptop^9^. NanoCortex covers nanopore analysis but emits generated code and is scored by a second language model rather than against ground truth^10^. BaseCamp configures short-read pipelines and reports its concordance as preliminary^11^. Seqera MCP launches Nextflow runs but publishes no biological validation^12^, and FlowBench, which does submit to schedulers, found that agents fail at fault recovery more than at planning^13^. None compares the output with an orthogonal measurement, and where a human reference exists the comparison is a rubric or similarity score, not identity of the output. Whether an agent can drive multi-hour GPU jobs on a scheduler and return the output a human expert would produce, verified against an orthogonal assay, has not been shown.

Long-read sequencing from Oxford Nanopore Technologies (ONT) and PacBio measures CpG methylation on native DNA, and because one read carries both the variants that define a haplotype and the methylation along it, phasing turns that measurement into allele-specific methylation, which imprinting, X inactivation and clonal analysis in cancer require^1,2^. The callers exist and are benchmarked on both platforms^3– 7^, and we previously assembled them into NANOME for nanopore data^8^. What still separates a biologist from the result is operation: a workflow manager, a cluster scheduler, GPU containers, and a basecalling model that has to match the chemistry of the run.

Four things have to be right before the first base is called, and none of them is visible in the answer if they are wrong. The basecalling model has to match the chemistry. PacBio is not the same route with a flag changed, because 5mC is read from polymerase kinetics that most public releases no longer carry. Allele-specific methylation needs variants, phasing and haplotagging to happen in order. And the work runs for hours on scheduled GPUs, where a failed task does not stop the run, so completion has to be read from the execution trace rather than from an exit code.

LongVerse (**Fig. 1a-c**) has three parts: a Nextflow^14^ pipeline, an MCP server that exposes it as 17 tools, and the agent that operates them. The pipeline runs from from raw ONT or PacBio signal to haplotype-resolved methylation, with one process per tool and every tool in a pinned container. ONT reads are basecalled and 5mC-called with Dorado; PacBio HiFi reads are 5mC-called from kinetics with Jasmine and aligned with pbmm2. Both branches then share quality control, variant calling (Sniffles2, Clair3)^15,16^, phasing (WhatsHap)^17^, per-read extraction of CpG calls, and differential methylation (methylKit, DSS)^18,19^, on T2T-CHM13v2.0^20^. The MCP server was built with Paper2Agent^9^ (**Fig. 1b**; **Supplementary Table S1**). Three rules make the agent verifiable: the tools read the run’s own data before a model is chosen, so chemistry, model and reference are read before any compute is spent; the agent supplies arguments and never code, so every computation stays inside the pinned pipeline; and the agent acts only at decision points, leaving the hours in between to the scheduler.

**Fig. 1.**
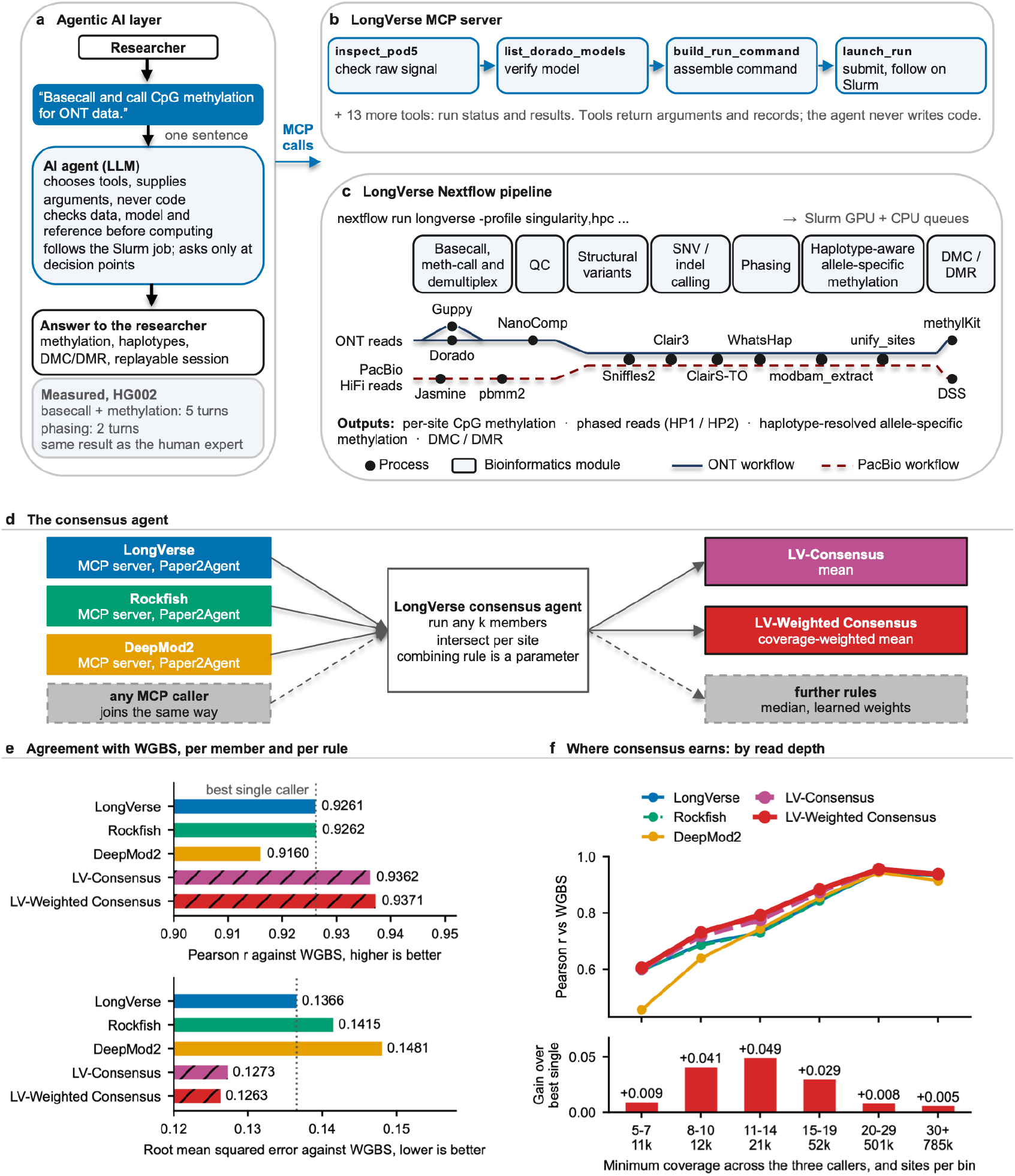
LongVerse: an agent layer, an MCP server, a pipeline, and consensus agents. **a**, The researcher states the task in one sentence; the agent chooses tools and supplies arguments, never code, checks data, model and reference before computing, and follows the Slurm job. **b**, The MCP server, built with Paper2Agent^9^, exposes the pipeline as 17 validated tools; for basecalling the agent calls inspect_pod5, list_dorado_models, build_run_command and launch_run. **c**, The Nextflow pipeline: ONT reads through Dorado, PacBio HiFi reads through Jasmine and pbmm2, then shared steps for quality control, variants, phasing, haplotype-aware methylation and differential methylation. **d**, The consensus agent: three members built with Paper2Agent (LongVerse, DeepMod2, Rockfish) and an open slot for any MCP caller; per-site calls are intersected and combined by a rule given as a parameter. **e**, Agreement with WGBS on 1,381,961 HG002 chromosome 22 CpG sites at coverage of five or more (Pearson r and RMSE), for the three members (LongVerse denotes the pipeline’s Dorado calls), the unweighted mean of all three (LV-Consensus) and the coverage-weighted mean (LV-Weighted Consensus). **f**, The same within six minimum-coverage bins, 5-7, 8-10, 11-14, 15-19, 20-29 and 30 or more reads, with the gain of LV-Weighted Consensus over the best single caller in each bin.

We then asked whether an agent, given one sentence, reproduces what an expert produces at the command line, each task run once per arm (**Fig. 2** and **Extended Data Fig. 1**; **Supplementary Tables S2-S4**). Given ‘Basecall and call CpG methylation for ONT data.’ and the raw signal of chromosome 20 from HG002, a healthy Genome in a Bottle cell line^21^, the agent inspected the POD5 files, listed the Dorado models in the container, composed the command and submitted it: five turns, 140 s, $0.54. Both arms completed all eight tasks; their per-site tables were identical (r = 1.000 over 1,575,278 CpGs) and each agreed with whole-genome bisulfite sequencing (WGBS) at r = 0.943. The expert’s command cost nothing but had to be right in eleven options, three of which fail silently while Nextflow exits 0. Given ‘Phase the reads and give me methylation per haplotype.’, the agent needed two turns and 62 s ($0.22); both arms completed 14 of 14 tasks with identical haplotype read sets and the same methylation at the chromosome 20 imprinting control regions (r = 1.000), down to region plots that match byte for byte.

**Fig. 2.**
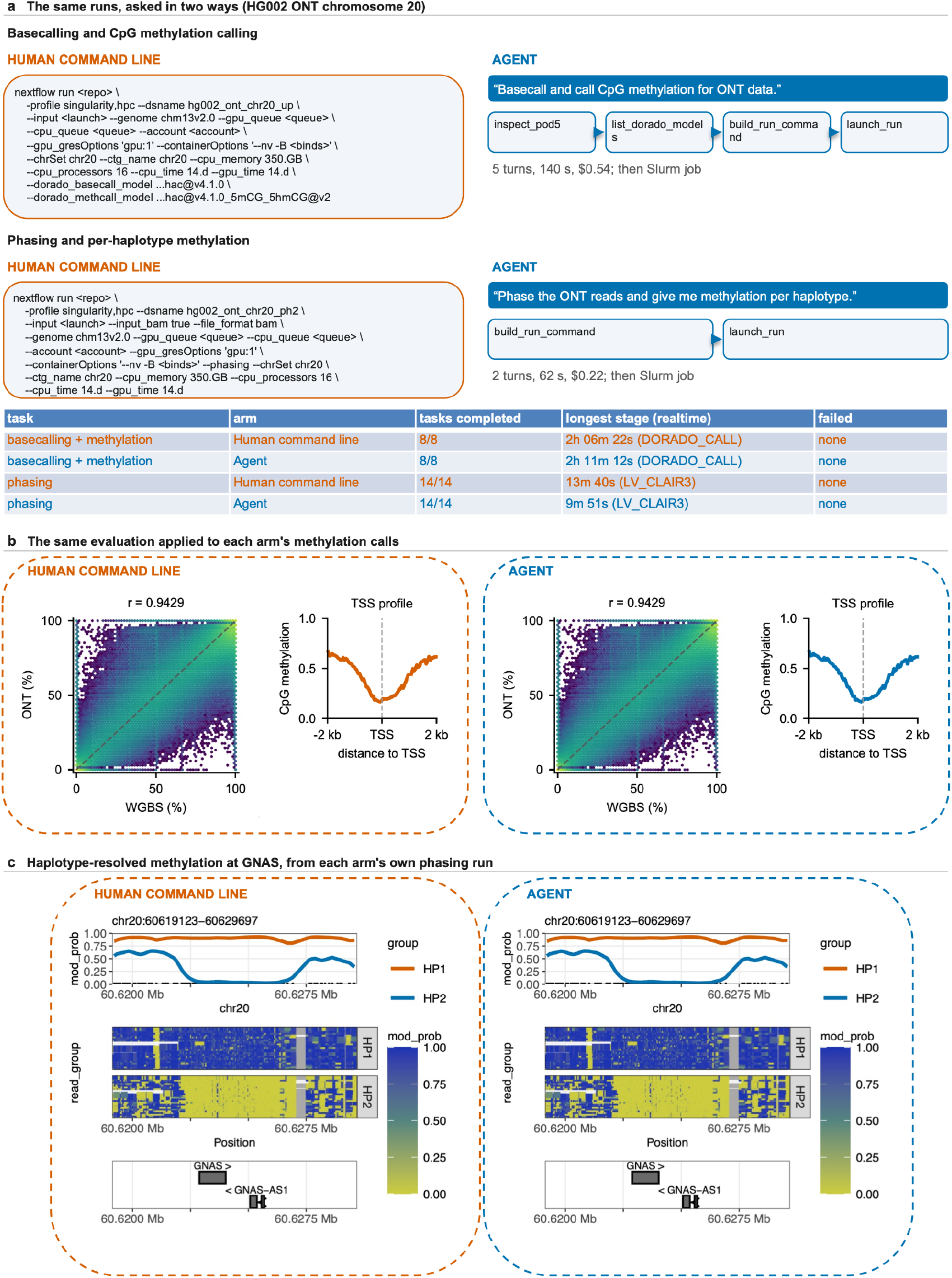
One sentence reproduces the expert’s methylation calling and phasing. **a**, Two runs on HG002 ONT chromosome 20, each asked in two ways: the expert’s nextflow command (abbreviated) and the agent’s sentence with the tools it chose, in the order it called them; tasks completed, longest stage and failures per arm, with the agent’s turns, tokens and cost. Wall time is conversation time and excludes the pipeline run it sets off. **b**, Each arm’s own methylation calls against WGBS (long reads at coverage of five or more) and as mean methylation in 50 bp bins over 2 kb each side of transcription start sites; the dashed frame groups the panels of one arm and carries its colour, orange for the expert at the command line and blue for the agent, throughout the figure. **c**, The GNAS imprinting cluster from each arm’s own phasing run (HP1 and HP2 methylation, read heatmap, gene track); the two panels are byte-identical, equal in md5 and not merely alike to the eye. The chromosome 20 imprinting control regions are in Extended Data Fig. 1.

The same held on PacBio HiFi kinetics for chromosome 20 (**Extended Data Figs. 2 and 3**; **Supplementary Tables S5 and S6**): all tasks completed, tables identical between runs, r = 0.884 against WGBS, $0.20 and $0.12 of conversation. The pipeline underneath scales to whole genomes, where the two platforms agree (**Extended Data Figs. 4 and 5**; **Supplementary Tables S7-S9**): 62.1 million CpG sites correlated at r = 0.882 between ONT and PacBio, and at r = 0.953 and 0.891 with WGBS, with the three assays in phase around transcription start sites and CTCF sites. Phasing gave 45,329 (ONT) and 21,570 (PacBio) haplotype-specific differentially methylated regions across the autosomes, and at the known imprinting control regions^1,2^ the haplotype difference was larger on ONT than on PacBio (38 regions, median 0.71 against 0.60, paired Wilcoxon P = 1.1 x 10^-7^).

Finally we asked whether agents built from different callers can be combined (**Fig. 1d-f**; **Supplementary Tables S10 and S11**). We built agents for two published nanopore callers, DeepMod2 and Rockfish^5,6^, and a consensus agent that contracts with the protocol: any MCP server returning per-site calls can join, and the combining rule is a parameter. Combining callers is established practice: METEORE^22^ improved accuracy by combining nanopore callers, as did NANOME^8^; what the consensus agent adds is the contract that lets any MCP caller join. We scored it on chromosome 22, because DeepMod2’s weights for this chemistry were trained on chromosomes 2 to 21 of this trio and chromosome 20 would score a model on its own training data. On 1,381,961 chromosome 22 CpG sites shared by all three callers and the bisulfite reference, the LongVerse pipeline’s Dorado calls and Rockfish reached r = 0.926 and DeepMod2 0.916; their coverage-weighted mean (LV-Weighted Consensus, **Fig. 1e,f**) reached r = 0.937 and a root mean squared error of 0.1263, 10.7% below the best published caller (Rockfish, r = 0.9262, error 0.1415) and below the pipeline’s own 0.1366, with the largest gains at 8 to 19x. The ceiling is the correlation of the callers’ errors, 0.66 to 0.81, because all three read the same basecalls.

Two boundaries are worth stating. Rockfish has one published checkpoint for this chemistry and it was trained at a different sampling rate, so every series containing it measures what a user would get today rather than a recommended configuration. And for the PacBio tasks the headless interface on this cluster refuses tool execution, so the agent chose the tool and its arguments and a scripted client ran the composed command; the cost reported there is the cost of deciding, which is the part that differs between an agent and an expert.

The agent’s contribution is not a different answer but the right choices: it read the run’s chemistry before picking a model, wrote no code, and left the computation to the pinned pipeline, which is why its output and the expert’s are byte-identical. That identity is what an agent layer should deliver, and what repository examples and language-model judges cannot certify. The tools report rather than decide: all five deliberately broken inputs composed into valid commands, and the only repair, a missing container bind, was reported as an addition rather than a fault (**Supplementary Table S12**). Fault recovery^13^ and more consensus members remain open, and the consensus contract is written so that the next member costs a registry entry rather than a rewrite. LongVerse gives a biologist haplotype-resolved methylation from one sentence, through checks it makes explicit and callers it can combine.

## Methods

### Data preprocessing

HG002 (GM24385) was analyzed throughout against the T2T-CHM13v2.0 reference^20^. For ONT, data from two PromethION flow cells (R10.4.1, LSK114, 4 kHz) were converted from FAST5 to POD5 and basecalled with Dorado 0.9.5 (dna_r10.4.1_e8.2_400bps_hac and 5mCG_5hmCG modification model), yielding approximately 68× aligned coverage. For PacBio, subreads from four Sequel II were reprocessed with ccs 6.4.0 --hifi-kinetics to retain kinetics tags. The resulting HiFi reads were merged to approximately 54× coverage; Jasmine called 5mC from the kinetics, and pbmm2 aligned the reads. For WGBS, GM24385 Swift Accel-NGS Methyl-Seq reads were trimmed with Trim Galore (15 bp from the 3′ end of read 1 and the 5′ end of read 2), aligned with Bismark and Bowtie2, deduplicated, and summarized per CpG with bismark_methylation_extractor and bismark2bedGraph --CX. Calls were used without filtering or strand merging. For the agent benchmark, chromosome 20 subsets of ONT signal and PacBio HiFi reads, plus a chromosome 22 ONT subset for consensus, were prepared so each run completed within hours.

### Pipeline

LongVerse is written in Nextflow^14^, with one process per tool. Dorado (GitHub: nanoporetech/dorado) uses basecalling and modification models on ONT data; Jasmine (GitHub: staphb/pbjasmine) calls 5mC from PacBio HiFi data. Other tools are pbmm2 (GitHub: PacificBiosciences/pbmm2), NanoComp (GitHub: wdecoster/nanocomp), Sniffles2^15^, Clair3 (GitHub: hkubal/clair3)^16^, and ClairS-TO (GitHub: HKU-BAL/ClairS-TO)^23^. WhatsHap^17^ performs haplotagging and splitting from the Clair3. Per-read CpGs are extracted from modified-base BAMs with canonical and modified probability thresholds of 0.5. methylKit^18^ and DSS^19^ are used for differential methylation.

### Phasing and allele-specific methylation

Clair3 is run with --enable_phasing and -- var_pct_phasing 0.80; variants with QUAL of at least 20 are kept; WhatsHap haplotag assigns reads to HP1 and HP2 and split writes one BAM per haplotype. Haplotype-specific differentially methylated cytosines and regions were called with methylKit and DSS; **Extended Data Fig. 4d,e** draws methylKit regions with q <= 0.01 and a difference of at least 50%. Imprinting control regions are the 91 intervals of Court et al.^1^ and Joshi et al.^2^ as compiled for NanoMethPhase^24^, lifted from hg38 to T2T-CHM13 with UCSC liftOver; 90 had coverage of five or more on both platforms in the region table, 43 novel and 38 known in the heatmaps. Per-region HP1/HP2 methylation was visualized by NanoMethViz^25^. The absolute difference between HP1 and HP2 mean methylation was compared between platforms per region set with a paired two-sided Wilcoxon signed-rank test.

### MCP server and agents

The LongVerse MCP server was built with Paper2Agent^9^ and lives in the pipeline repository (agents/longverse-mcp); its 17 tools are listed in **Supplementary Table S1**. The agent was Claude Opus 5 through the Claude Code command-line interface with only the LongVerse MCP tools attached. Two further MCP servers wrap DeepMod2^5^ and Rockfish^6^ (agents/consensus-mcp); the consensus agent takes any number of per-site callers, intersects sites and applies a combining rule given as a parameter (mean, coverage-weighted mean).

### Agent-versus-expert benchmark

For each task the expert typed one nextflow command and the agent received one sentence including the location of the data and reference. Both tests ran the same pipeline on the same HG002 chromosome 20 input and produced their own outputs. Evaluation was then applied identically to both outputs: Pearson correlation of per-site methylation with WGBS at coverage ≥ 5; correlation with WGBS by read-depth bin; mean methylation in 50 bp bins over +/-2 kb of RefSeq transcription start sites (hs1); HP1 and HP2 mean methylation over the chromosome 20 imprinting control regions; and NanoMethViz region plots.

### Consensus

On HG002 ONT chromosome 22, LongVerse, DeepMod2 and Rockfish were run through their MCP servers and their per-site tables intersected with the WGBS reference (1,381,961 sites at coverage ≥ 5). Each member, the mean of all three, the coverage-weighted mean were scored by Pearson r and RMSE against WGBS, overall and by minimum-coverage bin; residual correlations between members were computed to bound the attainable gain.

### Reporting Summary

Further information on research design is available in the Nature Portfolio Reporting Summary linked to this article.

## Data availability

HG002 ONT raw signal is from ONT open data (s3://ont-open-data/giab_lsk114_2022.12/flowcells/hg002/); PacBio subreads with kinetics are the DeepConsensus publication data^26^ (gs://brain-genomics-public/research/deepconsensus/publication/sequencing); HG002 (GM24385) bisulfite reads are from ONT open data (s3://ont-open-data/gm24385_mod_2021.09/bisulphite/).

## Code availability

LongVerse, its MCP server and the DeepMod2, Rockfish and consensus agents are at https://github.com/LabShengLi/longverse (public, release v0.0.1).

## Acknowledgements

This work was supported by NIH grants R35GM162228, R35GM133562, U01HG013175 and U01CA271830. Computation used the USC Center for Advanced Research Computing.

## Author contributions

Y.L. designed the pipeline and the agents, ran the benchmarks and wrote the manuscript; Q.Z. contributed the test code and wrote the manuscript; S.L. supervised the work and wrote the manuscript with input from all authors.

## Supplementary Tables

Table S1. The LongVerse MCP servers: every tool, the server it belongs to, and what it does Table S2. Per-site agreement with whole-genome bisulfite sequencing, ONT arms

Table S3. Haplotype-resolved methylation at chromosome 20 imprinting control regions, ONT arms

Table S4. Agreement with the bisulfite reference by read depth, ONT arms

Table S5. Per-site agreement with whole-genome bisulfite sequencing, PacBio arms Table S6. The two PacBio phasing arms compared per haplotype

Table S7. The three pairwise per-site comparisons as drawn, with coverage of five or more on both members

Table S8. Haplotype methylation at 90 imprinting control regions on ONT, PacBio (Jasmine) and a trio-phased reference

Table S9. ONT against PacBio per-site methylation within genomic regions

Table S10. Three callers and their consensus against whole-genome bisulfite sequencing Table S11. The same comparison split by read depth

Table S12. Five deliberately broken inputs through the command composer

**Extended Data Fig. 1.**
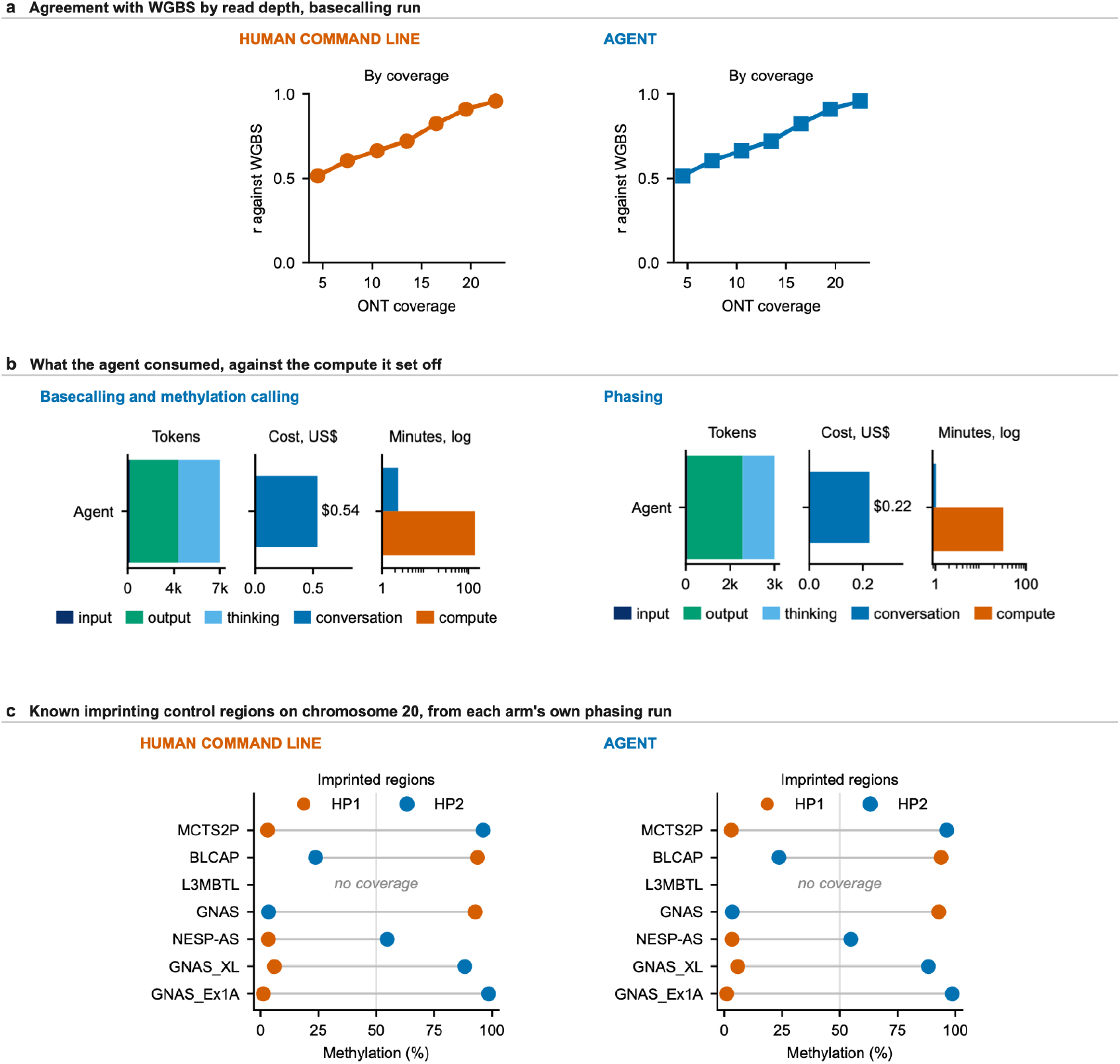
The ONT agent-versus-expert comparison. **a**, Agreement with WGBS by read-depth bin for each arm (basecalling run). **b**, The agent’s tokens, cost and conversation time against the compute each run set off, for the basecalling and the phasing task. **c**, The known chromosome 20 imprinting control regions as per-haplotype methylation, from each arm’s own phasing run and byte-identical between them (md5 stated); both runs completed 14 of 14 tasks and assigned 95,576 and 95,190 reads to the two haplotypes with identical read sets.

**Extended Data Fig. 2.**
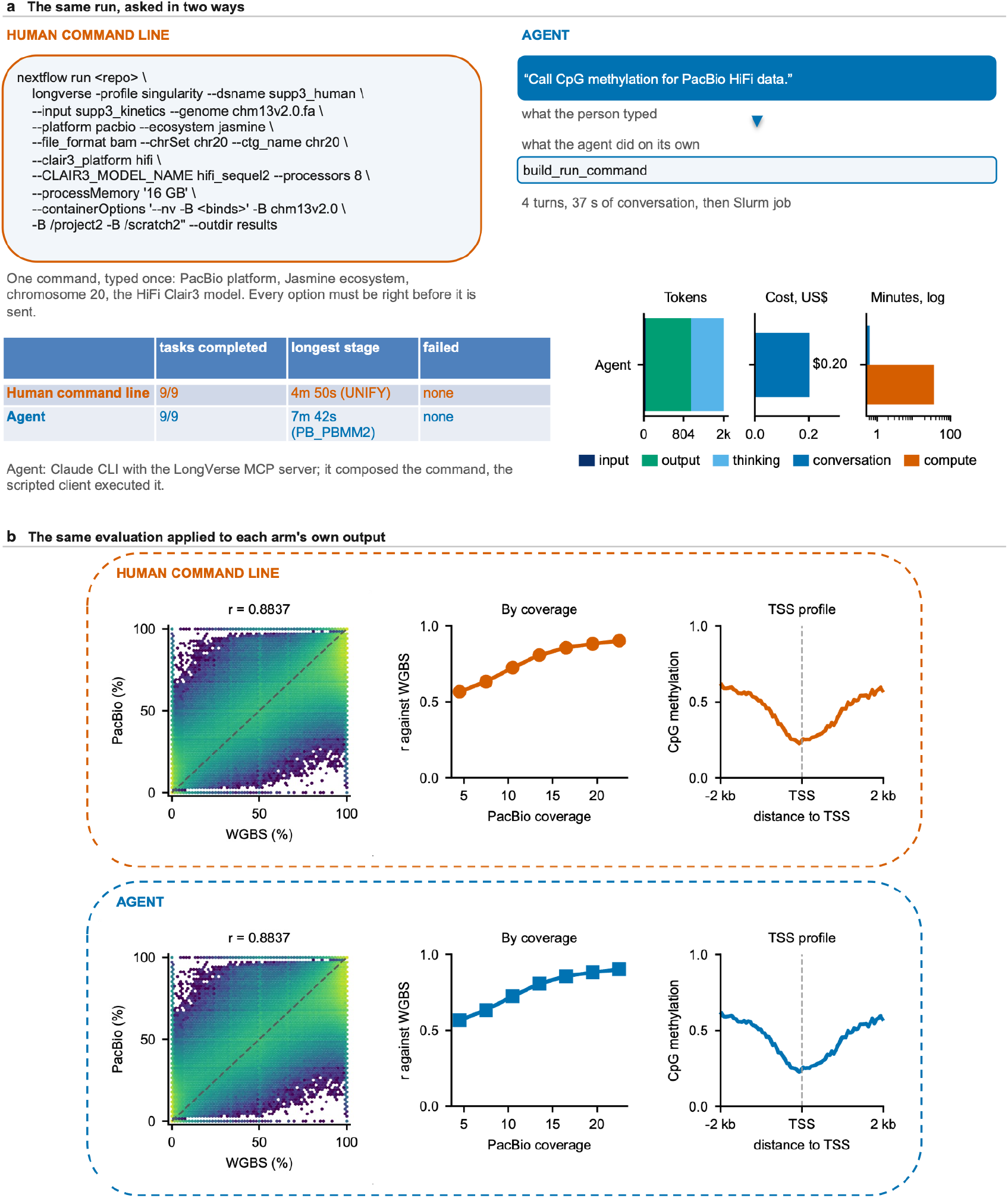
One sentence reproduces a PacBio methylation-calling run. **a**, The same run asked in two ways, as Fig. 2a, for HG002 PacBio HiFi reads with kinetics on chromosome 20 (Jasmine 5mC, pbmm2): the expert’s nextflow command and the agent’s sentence with the tool it chose; tasks completed, longest stage and failures per arm, with the agent’s tokens, cost and conversation time (4 turns, 37 s, $0.20). **b**, The same evaluation applied to each arm’s own output, as Fig. 2b: per-site methylation against WGBS (r = 0.884 over 1,541,666 CpGs), agreement with WGBS by read-depth bin and the TSS profile; 9 of 9 tasks in both runs, per-site tables identical (r = 1.000 over 1,574,491 sites).

**Extended Data Fig. 3.**
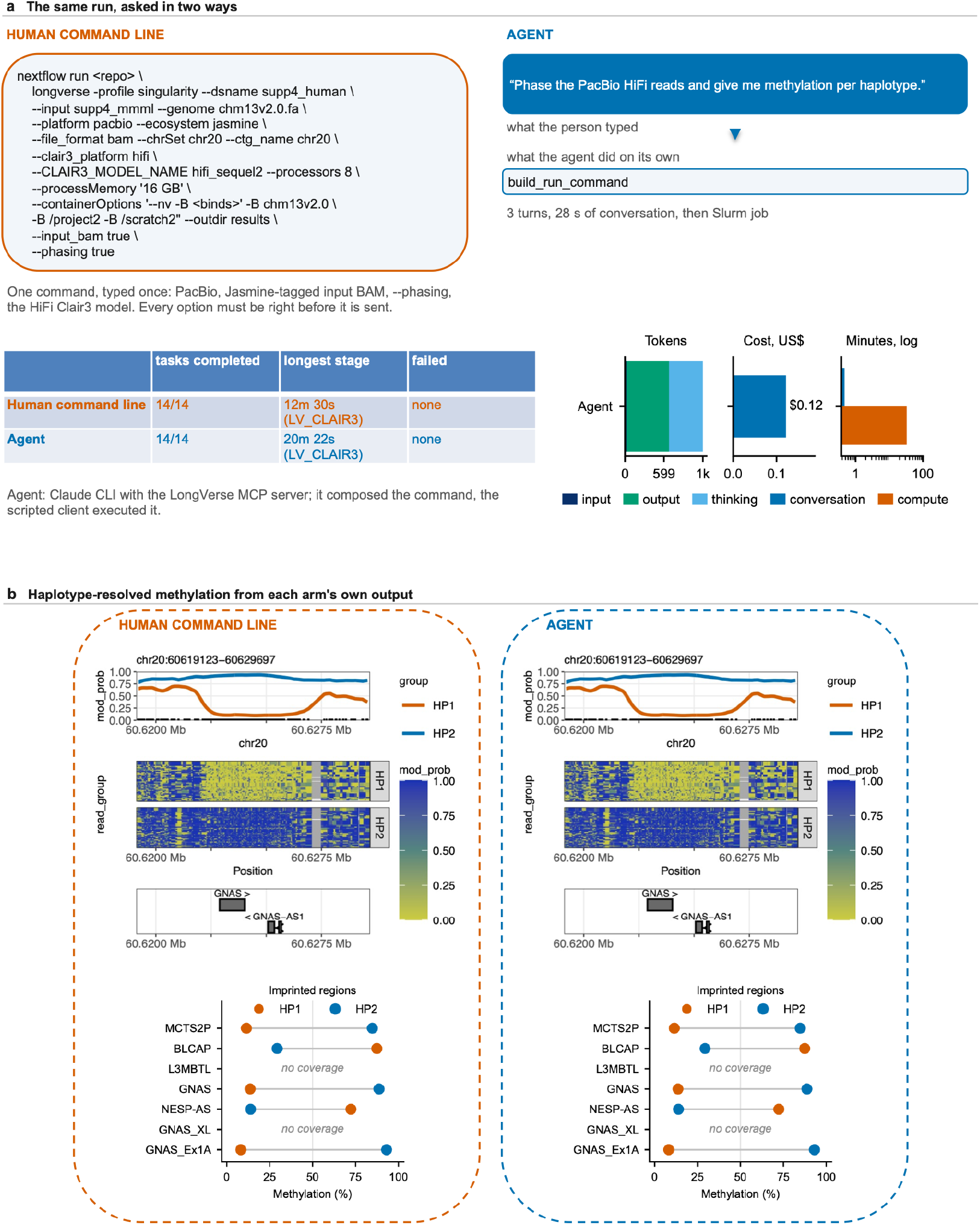
One sentence reproduces a PacBio phasing run. **a**, The same run asked in two ways, as Fig. 2a, for the same PacBio reads carrying 5mC tags: the expert’s nextflow command with phasing and the agent’s sentence with the tool it chose; tasks completed, longest stage and failures per arm, with the agent’s tokens, cost and conversation time (3 turns, 28 s, $0.12). **b**, Haplotype-resolved methylation from each arm’s own output, as Fig. 2c: the GNAS imprinting cluster (HP1 and HP2 methylation, read heatmap, gene track) and the known chromosome 20 imprinting control regions per haplotype; 14 of 14 tasks in both runs, HP1 and HP2 per-site tables identical (1,385,404 and 1,380,334 sites, r = 1.000).

**Extended Data Fig. 4.**
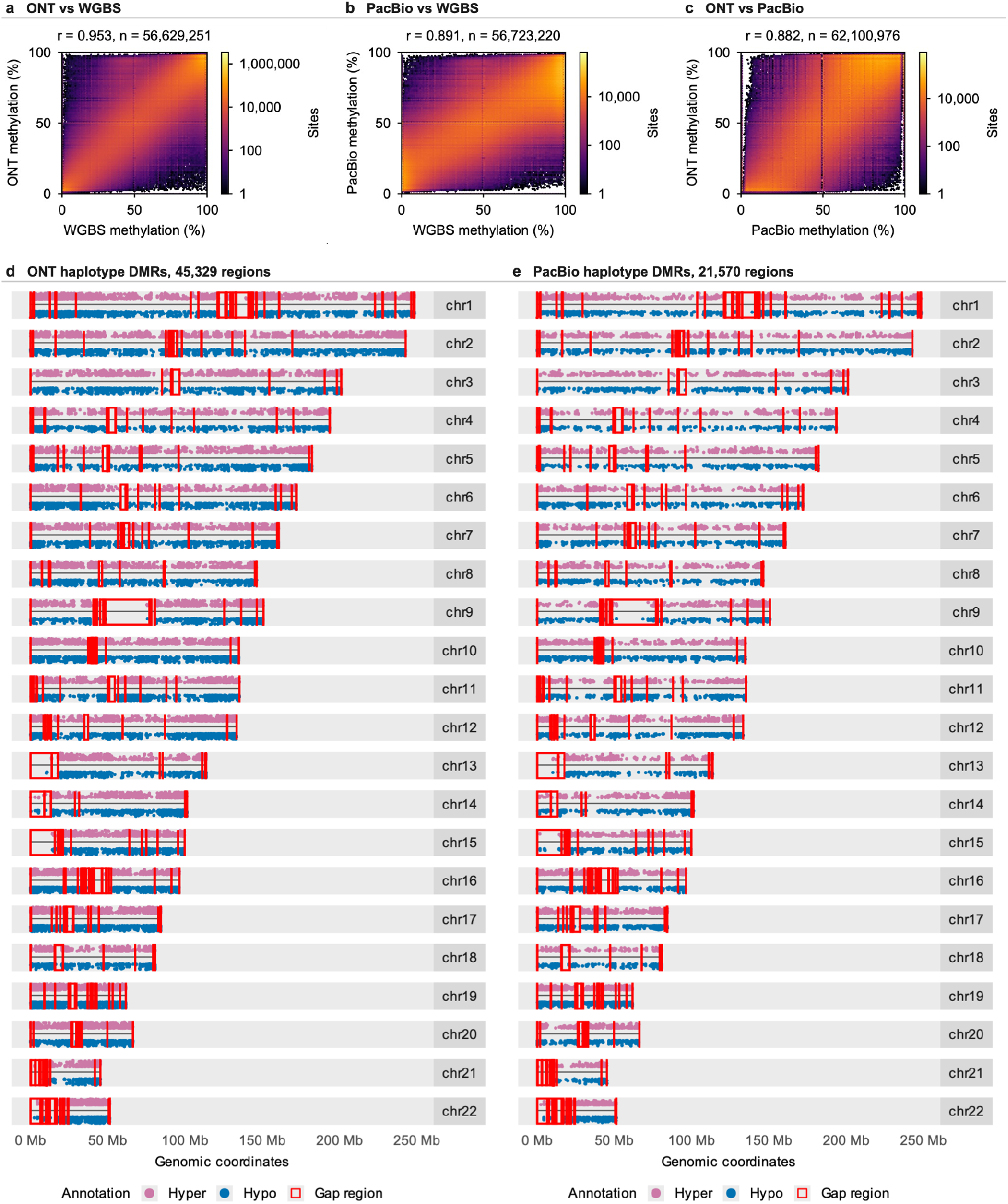
LongVerse on ONT and PacBio, HG002 whole genome, coverage of five or more on both platforms. **a**, Per-site methylation, ONT against WGBS (56,629,251 CpG sites, r = 0.953). **b**, PacBio against WGBS (56,723,220 sites, r = 0.891). **c**, ONT against PacBio (62,100,976 sites, r = 0.882). **d**,**e**, Haplotype-specific differentially methylated regions (methylKit, HP1 against HP2, 100 bp tiles, q <= 0.01, difference >= 50%) on every autosome for ONT (d; 45,329 regions) and PacBio (e; 21,570); hypermethylated regions in pink, hypomethylated in blue, T2T-CHM13 gap regions outlined in red. PacBio 5mC was called from HiFi kinetics inside the pipeline (Methods).

**Extended Data Fig. 5.**
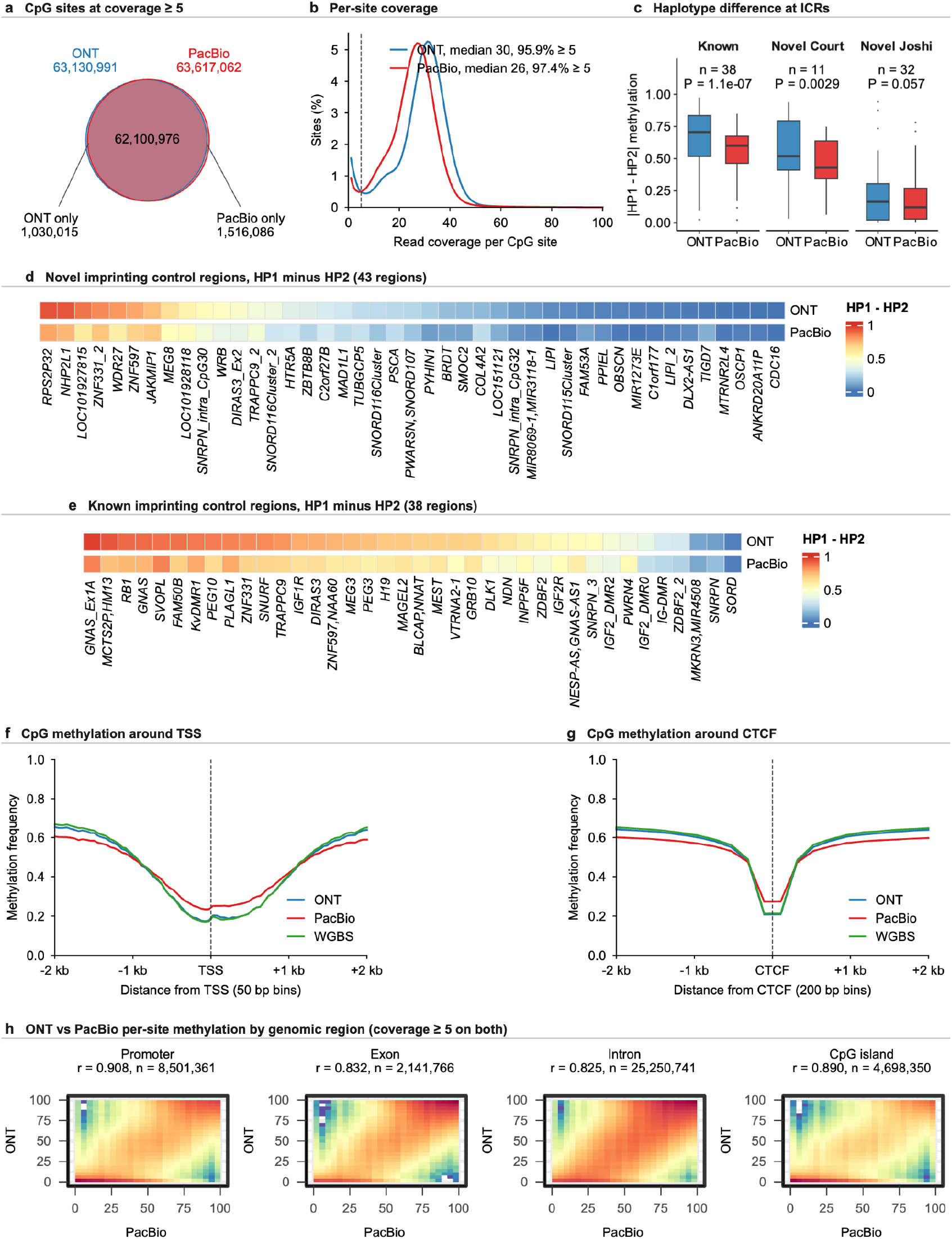
Coverage, imprinting control regions and regional agreement of the two platforms, HG002 whole genome. **a**, CpG sites covered at five or more reads by ONT (63,130,991), by PacBio (63,617,062) and by both (62,100,976). **b**, Per-site coverage distributions for all called sites (ONT 65,801,544 sites, median 30, 95.9% at five or more; PacBio 65,314,640, median 26, 97.4%); the dashed line marks coverage five. **c**, Absolute haplotype difference at the 81 imprinting control regions with a haplotype difference on both platforms, by region set (known 38, novel Court 11, novel Joshi 32 regions); P values from paired two-sided Wilcoxon signed-rank tests. **d**,**e**, HP1 minus HP2 methylation at the regions reported as novel (d; 43 regions) and known (e; 38 regions), ONT above PacBio. **f**,**g**, Mean CpG methylation around transcription start sites (f) and CTCF sites (g) for ONT, PacBio and WGBS, over 2 kb each side; bins of 50 bp (f) and 200 bp (g), each averaging the CpG sites it contains, coverage of five or more on every track. The wider bin in g averages over the nucleosome-phased oscillation that the three assays show in phase around CTCF sites. **h**, ONT against PacBio per-site methylation within four genomic regions (promoter, exon, intron, CpG island), colour showing the number of sites in each two-dimensional bin, with Pearson r and the total above each panel.

**Extended Data Table 1.** Agentic and pipeline systems for sequencing analysis.

| System | Scope | Cluster jobs | Checks data first | Agent output | Validated against | Combines callers |
| --- | --- | --- | --- | --- | --- | --- |
| Paper2Agent | Any paper to an MCP server and agent; laptop-scale tools (22 AlphaGenome tools in 45 min for \$14) | No | No | Tool calls | Human-executed ground truth, graded by two experts: 98.7% tutorial, 82.7% open-ended queries | No |
| NanoCortex | Nanopore, basecalling to interpretation; RNA modification, isoform and fusion tasks | No (local) | Reads the user's data description | Generated code (Python, R, Bash) | Capability rubric scored by a language model over documented features; no ground truth | No |
| BaseCamp | Short-read DNA: QC, alignment, variant calling, annotation; six fine-tuned local agents | No (on-premises) | Yes (QC reports, alignment statistics) | Tool selection and configuration | Expert manual configurations (concordance; preliminary) | No |
| FlowBench / FlowAgent | Benchmark: planning, fault recovery, interpretation; 23 models | Yes (SLURM, SGE, TORQUE via DRMAA) | n/a | Workflow plan (JSON DAG) | Pre-registered toolchains; automated scoring; no human baseline | n/a |
| Seqera MCP | Launch, monitor and manage Nextflow runs on Seqera Platform | Via Seqera compute environments | No | Workflow launch | None published | No |
| **LongVerse (this work)** | ONT and PacBio, raw signal to allele-specific methylation; 17 MCP tools; consensus agent | Yes (Slurm, asynchronous) | Yes (inspect_pod5, list_dorado_models) | Arguments only, never code | WGBS and a human expert (byte-identical outputs) | Any MCP caller |

## References

1. Court, F. et al. Genome-wide parent-of-origin DNA methylation analysis reveals the intricacies of human imprinting and suggests a germline methylation-independent mechanism of establishment. Genome Res. 24, 554–569 (2014). PMID 24402520.

2. Joshi, R. S. et al. DNA methylation profiling of uniparental disomy subjects provides a map of parental epigenetic bias in the human genome. Am. J. Hum. Genet. 99, 555–566 (2016). PMID 27569549.

3. Sigurpalsdottir, B. D. et al. A comparison of methods for detecting DNA methylation from long-read sequencing of human genomes. Genome Biol. 25, 69 (2024). PMID 38468278.

4. Kulkarni, O. et al. Comprehensive benchmarking of tools for nanopore-based detection of DNA methylation. Nat. Commun. 17, 8685 (2026). PMID 42457719.

5. Ahsan, M. U., Gouru, A., Chan, J., Zhou, W. & Wang, K. A signal processing and deep learning framework for methylation detection using Oxford Nanopore sequencing. Nat. Commun. 15, 1448 (2024). PMID 38365920.

6. Stanojevic, D., Li, Z., Bakic, S., Foo, R. & Sikic, M. Rockfish: a transformer-based model for accurate 5-methylcytosine prediction from nanopore sequencing. Nat. Commun. 15, 5580 (2024). PMID 38961062.

7. Ni, P. et al. DNA 5-methylcytosine detection and methylation phasing using PacBio circular consensus sequencing. Nat. Commun. 14, 4054 (2023). PMID 37422489.

8. Liu, Y. et al. NANOME: a Nextflow pipeline for haplotype-aware allele-specific consensus DNA methylation detection by nanopore long-read sequencing. Preprint at bioRxiv (2025). PMID 40631091, PMC12236756.

9. Miao, J., Davis, J. R., Zhang, Y., Pritchard, J. K. & Zou, J. Reimagining research papers as interactive and reliable AI agents. Nature 10.1038/s41586-026-11044-y (2026). PMID 42749808.

10. Xia, Q., Wang, Z., Shokoufandeh, M., Rouhanifard, S. H. & Wanunu, M. NanoCortex: a unified agentic system for nanopore sequencing analysis. Preprint at bioRxiv 10.64898/2026.05.19.726254 (2026). PMID 42239323.

11. Bandara, E. et al. BaseCamp: an agentic AI framework for automating DNA sequencing data pipelines. Preprint at arXiv https://arxiv.org/abs/2609.28557 (2026).

12. Seqera Labs. Seqera MCP. https://docs.seqera.io/platform-cloud/seqera-mcp (2025).

13. Kurjan, A. & Cribbs, A. P. FlowBench: separating planning, fault recovery and interpretation in agentic bioinformatics. Preprint at bioRxiv 10.64898/2026.06.12.731844 (2026).

14. Di Tommaso, P. et al. Nextflow enables reproducible computational workflows. Nat. Biotechnol. 35, 316–319 (2017). PMID 28398311.

15. Smolka, M. et al. Detection of mosaic and population-level structural variants with Sniffles2. Nat. Biotechnol. 42, 1571–1580 (2024). PMID 38168980.

16. Zheng, Z. et al. Symphonizing pileup and full-alignment for deep learning-based long-read variant calling. Nat. Comput. Sci. 2, 797–803 (2022). PMID 38177392.

17. Patterson, M. et al. WhatsHap: weighted haplotype assembly for future-generation sequencing reads. J. Comput. Biol. 22, 498–509 (2015). PMID 25658651.

18. Akalin, A. et al. methylKit: a comprehensive R package for the analysis of genome-wide DNA methylation profiles. Genome Biol. 13, R87 (2012). PMID 23034086.

19. Feng, H., Conneely, K. N. & Wu, H. A Bayesian hierarchical model to detect differentially methylated loci from single nucleotide resolution sequencing data. Nucleic Acids Res. 42, e69 (2014). PMID 24561809.

20. Nurk, S. et al. The complete sequence of a human genome. Science 376, 44–53 (2022). PMID 35357919.

21. Zook, J. M. et al. Extensive sequencing of seven human genomes to characterize benchmark reference materials. Sci. Data 3, 160025 (2016). PMID 27271295.

22. Yuen, Z. W.-S. et al. Systematic benchmarking of tools for CpG methylation detection from nanopore sequencing. Nat. Commun. 12, 3438 (2021). PMID 34103501.

23. Chen, L. et al. ClairS-TO: a deep-learning method for long-read tumor-only somatic small variant calling. Nat. Commun. 16, 9630 (2025). PMID 41173866.

24. Akbari, V. et al. Megabase-scale methylation phasing using nanopore long reads and NanoMethPhase. Genome Biol. 22, 68 (2021). PMID 33618748.

25. Su, S. et al. NanoMethViz: an R/Bioconductor package for visualizing long-read methylation data. PLoS Comput. Biol. 17, e1009524 (2021). PMID 34695109.

26. Baid, G. et al. DeepConsensus improves the accuracy of sequences with a gap-aware sequence transformer. Nat. Biotechnol. 41, 232–238 (2023). PMID 36050551.

